# Intermittent exposure to high ambient heat during the second half of gestation in mice causes mild alterations of reproductive endpoints in male embryos

**DOI:** 10.64898/2026.05.22.727256

**Authors:** Kimberly Abt, Ciro M. Amato, Abigail Kitakule, Yu-Ying Chen, Barbara Nicol, Karina Rodriguez, Erixberto Olivencia Álvarez, Carlos Guardia, Sara Grimm, Leslie Aksu, Korey Stevanovic, Jesse Cushman, Humphrey Hung-Chang Yao

**Affiliations:** Reproductive Developmental Biology Group, National Institute of Environmental Health Sciences, Research Triangle Park, NC 27709, USA; Department of Surgery, Division of Urology, University of Missouri, Columbia, MO 65211, USA; Department of Biological Sciences, University of Missouri, Columbia, MO 65211, USA; NextGen Precision Health, University of Missouri, Columbia, MO 65211, USA; Integrative Bioinformatics Support Group, National Institute of Environmental Health Sciences, Research Triangle Park, NC 27709, USA; Neurobehavioral Core Facility, National Institute of Environmental Health Sciences, Research Triangle Park, NC 27709, USA

**Author notes:** Corresponding Author: Humphrey H-C Yao. Authors contributed equally (Kimberly Abt, Ciro M. Amato).

**Keywords:** Heat Stress, Pregnancy, Hypospadias, Fetal Testis

## Abstract

Periods of elevated ambient temperature challenge the body’s ability to maintain internal homeostasis, and heat stress poses particular risks during pregnancy. Epidemiological studies associate gestational heat exposure with higher rates of congenital anomalies such as hypospadias, yet the direct link between gestational heat exposure and reproductive anomalies remains unknown. In this study, we examined the effects of intermittent heat exposure on reproductive development in male mouse offspring. Pregnant dams either remained at constant temperature of 22°C (control) or were exposed to 38°C for 2 hours daily (experimental) from embryonic day (E)10 to E18, modeling intermittent heat exposure during mid-to-late gestation. Embryos were collected at E18 for analysis. While heat exposure did not affect pregnancy outcomes, including placental development, litter size, sex ratio, or fetal growth, male embryos exhibited significantly reduced anogenital distance and increased hypospadias scores, which are both markers of disrupted androgen signaling. Despite these phenotypic changes, expression of genes involved in androgen synthesis in the fetal testis, as well as gene expression in external genitalia, remained unchanged. Instead, transcriptomic analysis revealed significant alterations in testicular pathways related to RNA splicing and mRNA processing. Together, these findings reveal that maternal heat stress disrupts reproductive development of male offspring, with altered gene regulatory processes being a potential driver.

## 1. Introduction

Heat stress occurs when external temperatures exceed the body’s capacity to maintain internal thermal homeostasis, which triggers alterations in metabolism, endocrine signaling, and cardiovascular function [1, 2]. During pregnancy, maternal physiological responses to heat stress can influence the intrauterine environment that supports fetal growth and development [2]. Changes in maternal body temperature, circulating hormones, and metabolic status can affect placental function and fetal organogenesis [3]. Increasing evidence suggests that maternal exposure to elevated ambient temperatures during gestation is associated with adverse developmental outcomes in offspring, including congenital abnormalities and perinatal death [1, 4–6]. However, the mechanisms through which gestational heat exposure influences fetal development remain poorly understood.

One congenital abnormality of particular concern is hypospadias, the most common birth defect affecting the male reproductive tract, which occurs in approximately 1% of newborn boys [7–9]. Hypospadias result from incomplete closure of the urethral tube during development of the penis, causing the urethral opening to form along the ventral shaft rather than at the distal tip. Although hypospadias is relatively common, the causes of approximately 70% of cases remain unknown. Epidemiological studies in humans have suggested a positive association between gestational exposure to elevated ambient temperatures and hypospadias incidence; however, a causal relationship between maternal heat exposure and this condition has not been established [10].

Development of the male reproductive system during fetal life is a highly coordinated process regulated by endocrine signaling and cellular interactions. Following sex determination, the fetal testes differentiate and begin producing testosterone from Leydig cells, which drives masculinization of the reproductive tract and supports development of the external genitalia [11, 12]. Penile development begins with formation of the genital tubercle, which undergoes morphogenesis during mid to late gestation. Proper penis formation requires coordinated signaling between epithelial and mesenchymal cell populations and adequate androgen production, which together lead to closure of the urethral tube [13–16]. Disruptions in fetal testis development, androgen signaling, or cellular interactions during this critical developmental window can impair urethral closure and result in abnormalities such as hypospadias.

Previous studies have demonstrated that exposure of pregnant rodents to elevated ambient temperatures can impair fetal growth, reduce placental development, and increase developmental abnormalities [17–21]. For example, exposure of pregnant mice to temperatures above 42°C for one hour from embryonic day (E) 6.5 to E14.5 led to reductions in fetal weight, crown rump length, and placental growth [17]. The same exposure scheme during late gestation (E14.5 to E17.5) altered maternal physiology and reduces placental size, indicating that gestational heat exposure can disrupt the fetal environment that supports proper development [17]. Other studies in mice found that exposure to 37°C for eight hours from E15.5 to E17.5 resulted in reduced fetal and placental size [19]. Importantly, exposure to 34°C for eight hours per day from E8 to E18 resulted in altered reproductive endpoints in male offspring, including reduced anogenital distance at birth and decreased adult testis size in mice, while female anogenital distance was not affected [20]. Despite growing evidence in animal models that gestational heat exposure disrupts reproductive development of male offspring, the specific developmental windows during which it acts, the underlying molecular mechanisms involved, and the long-term reproductive consequences remain unclear.

In this study, we developed a heat exposure model to determine whether intermittent gestational heat stress affects reproductive development in male embryos in mice. Pregnant dams were exposed to controlled elevated temperatures during the critical developmental window for reproductive organ formation. In contrast to other studies, this model uses intermittent daily heat exposure to mimic short-term periods of high environmental temperature that may occur during the day, rather than continuous heat exposure. This design allows us to model repeated short-term heat events during pregnancy while maintaining normal temperatures during the remainder of the day. Using this approach, we investigated how gestational heat exposure affects developmental endpoints in male offspring, including fetal testis development and penile morphology. Establishing an experimental model of intermittent gestational heat stress will help clarify whether elevated temperature during pregnancy contributes to developmental abnormalities of the male reproductive tract and may provide insight into environmental risk factors for hypospadias.

## 2. Materials and Methods

### 2.1 Animals and Heat Exposure Regime

All experimental procedures were approved by the National Institute of Environmental Health Sciences ACUC, ASP# 2010-0016. All animals were randomly assigned to control or experimental groups. Female C57BL/6 mice, 9-18 weeks old (Jackson Laboratories, Bar Harbor, ME), were time mated with C57BL/6 males. The day a vaginal plug was observed was considered embryonic day (E) 0.5 and pregnant females were weighed and individually housed in plastic cages with Sani-Chips bedding (PJ Murphey Forest Products Corp). Mice were provided NIH-31 chow *ad libitum* throughout the course of the experiment. During intermittent heat exposure, hydrogel (Lab Supply, 70-01-1082) and wet chow were provided to both control and experimental groups to prevent dehydration. At E6.5 females were weighed and ultrasound imaging was performed using a Vevo 3100 imaging platform (FUJIFILM visual systems) to confirm pregnancy. If embryonic structures were detected via ultrasound, the female was considered pregnant and transferred to temperature-controlled, animal-rearing chambers (Caron Scientific, 7350-33) to allow for habitat acclimation at 22°C. Two separate heat exposures were conducted at E10. In the first experiment, four mice were housed in four separate cages and were designated as the control group (22°C). Twelve mice were designated as the experimental group exposed to 40°C. In the second experiment, six mice were housed within six separate cages in one Caron chamber and were designated as the control group (22°C). Ten mice were designated as the experimental group exposed to 38°C. All cages had cameras to allow for live viewing during experimental conditions.

### 2.2 Mouse dissections

At E18.5 all pregnant mice were euthanized via CO_2_ inhalation. Maternal blood was collected by cardiac puncture in Greiner blood collection system tubes (Fisher Scientific Cat # 22-030-401). Pregnant dams were then dissected. All embryos were removed from the uterus and weighed. Placentas were collected, weighed, and diameter was measured. Half of the placenta was preserved in 4% paraformaldehyde for histology and the other half was snap-frozen for RNA-sequencing. Anogenital distance of the embryos was measured with digital electronic calipers (Chicago Brand, Part number 50013) from the center of the anus to the base of the external genitalia. Embryos were fully decapitated and the body cavity opened. Gonads were collected to determine the sex of each embryo. One testis was collected in 4% paraformaldehyde for histological sectioning, and the other testis was snap frozen for mRNA sequencing. Male external genitalia were then dissected, imaged, and half of the samples preserved in 4% paraformaldehyde, and the other half snap frozen for mRNA analysis.

### 2.3 Identifying hypospadias severity

All male external genitalia were imaged with a Leica model MP641 stereomicroscope. To identify hypospadias the previously published protocol was used to score individual genitalia [22]. Briefly, images were de-identified, randomized, and scored by two independent researchers. A score of 0 indicates no hypospadias, a score of 1 indicates mild/distal hypospadias, a score of 2 indicates moderate/mid-shaft hypospadias, and a score of 3 indicates severe/proximal hypospadias. Scores were averaged between the two researchers and analyzed. Logistic regression was then used to statistically determine hypospadias incidence rates.

### 2.4 Immunofluorescence of fetal testes

Testes were fixed in 4% paraformaldehyde at 4°C overnight. Samples were then washed three times for 10 minutes with 1x phosphate buffered saline (PBS) at room temperature. Testes were stored at 4 °C in 70% ethanol and subsequently embedded in paraffin. Tissue sections (5 μm) were deparaffinized, rehydrated, and subjected to antigen retrieval using a 1:1000 dilution of citrate-based antigen unmasking solution (Vector Laboratories, H-3300-250). The solution was pre-heated in a microwave for 5 minutes at full power, and slides were then heated for 20 minutes at 10% power. After cooling at room temperature for 30 minutes, sections were transferred to a Sequenza manual immunohistochemistry system and processed in PBS (100 μL per step unless otherwise indicated). Sections were blocked and permeabilized for 1 hour in buffer consisting of 0.1% Triton X-100 in 1x PBS supplemented with 5% normal donkey serum. Primary antibodies against GCNA (DiagnoCine,73-003), NR2F2 (R&D Systems, PPH7147-00), and 3β-HSD (CosmoBio, K0607) were diluted 1:200 in blocking buffer and applied overnight at 4°C. Sections were washed three times with 0.1% Triton X-100 in 1x PBS and incubated with secondary antibodies (Alexa Fluor 488, 594, and 647) diluted in blocking buffer for 1 hour at room temperature. Slides were washed once with 0.1% Triton X-100 in 1x PBS and twice with 1x PBS, followed by treatment with the TrueView Autofluorescence Quenching Kit (Vector Laboratories, SP-8400) for 3 minutes. Sections were washed with 500 μL of 1x PBS, counterstained with DAPI (Invitrogen, D1306), rinsed once more in 1x PBS, and mounted using ProLong™ Diamond Antifade Mountant (Invitrogen, P36970). Images were acquired using a Zeiss LSM 900 confocal microscope with Zen software. Brightness and contrast were adjusted using FIJI.

### 2.5 Hormone measurements

All blood was set at room temperature for >30 minutes to allow for clotting. Samples were then spun at 8,000 RCF for eight minutes and serum was collected and stored at - 80°C until further processing. Samples were sent to Wisconsin National Primate Research Center for LC-MS/MS analysis. Corticosteroid, estradiol, progesterone, testosterone, and dihydrotestosterone were all analyzed within maternal serum.

### 2.6 RNA Extraction

RNA was extracted from the external genitalia of four male control embryos and ten male experimental embryos; from the testes of eight control embryos and eleven experimental embryos; and from the placentas of seven male control embryos and seven male experimental embryos. Total RNA was isolated using the Arcturus™ PicoPure™ RNA Isolation Kit (Applied Biosystems, KIT0204) according to the manufacturer’s instructions. Samples were homogenized in 100 μL extraction buffer using a microtube homogenizer and incubated at 42°C for 30 minutes, followed by storage at −80°C for a minimum of 1 hour. RNA was precipitated by adding 100 μL of 70% ethanol and applied to a pre-conditioned purification column. Following centrifugation, columns were washed with Wash Buffer 1 and treated with 40 μL RNase-Free DNase Set (Qiagen, #79254) to remove genomic DNA. Columns were subsequently washed twice with Wash Buffer 2 and centrifuged to remove residual wash buffer. RNA was eluted in 13 μL of elution buffer. RNA concentration was determined using the Qubit™ RNA High Sensitivity Assay Kit (Thermo Fisher Scientific, Q32852). RNA integrity was evaluated using RNA ScreenTape (Agilent Technologies, #5067-5576) on an Agilent 4200 TapeStation system.

### 2.7 Bulk mRNA sequencing

For external genitalia and placenta samples, 250 ng of RNA were used for libraries preparation using the TruSeq Stranded mRNA Library Prep protocol (Illumina, #20020594). For fetal testis samples, libraries were made using Tecan’s Ovation RNA-Seq System V2 followed by Tecan’s Celero™ EZ DNA-Seq. Libraries were prepared and sequenced by the NIEHS Epigenomics and DNA Sequencing Core as paired-end 151-mers on an Illumina NextSeq 500 instrument.

### 2.8 Analysis of mRNA sequencing data

Read pairs were mapped to the mm10 reference genome via STAR v2.5.1b with parameters “--outSAMattrIHstart 0 --outFilterType BySJout --alignSJoverhangMin 8 --limitBAMsortRAM 55000000000 --outSAMstrandField intronMotif --outFilterIntronMotifs RemoveNoncanonical”. Counts per gene were determined via featureCounts (Subread v1.5.0-p1) with parameters “-s0-Sfr-p” for the testis dataset or parameters “-s2-Sfr-p” for the placenta and external genitalia datasets. Evaluated gene models were taken from the NCBI RefSeq Curated annotations as downloaded from the UCSC Table Browser on April 21, 2021. DESeq2 v1.42.059 was used for principal component analysis and identification of differentially expressed genes (threshold set at FDR 0.05) [23]. Gene ontology and pathway analysis were analyzed using Enrichr [24] and graphed using R studio.

### 2.9 Statistical Analysis

For comparisons of litter size, fetal weight or length, placental morphology, serum hormone levels, anogenital distance, and hypospadias severity score an unpaired two-tailed t-test was used to calculate statistical significance (P < 0.05). Data visualization and statistical analyses were performed using GraphPad Prism.

## 3. Results

### 3.1 Design of an intermittent heat exposure model during the second half of gestation in mice

To mimic intermittent heat exposure that humans would experience during the day, we designed the following experimental model using the climate control environmental chamber (Figure 1A): pregnant mice were exposed to rising temperatures during their active period after the light was turned off at 18:00 (Figure 1A). For the experimental group, starting at 20:00, the temperature in the environmental chamber was gradually ramped up (one hour of time) from 22°C to 38°C or 40°C, remained at 38°C or 40°C for 2 hours, and ramped down to 22°C (one hour). During the same time, pregnant mice in the control group remained at a constant 22°C. Hydrogel and wet chow were provided to both control and experimental groups to prevent dehydration. To determine if the mice suffered from significant dehydration, both control and experimental pregnant dams were weighed immediately prior to and after the experimental procedures. The same experimental procedure was conducted daily during the critical window of male sex differentiation, E10.0-E18.0 (Figure 1A). In the first attempt, we exposed the pregnant mice to 40°C based on heatwave temperatures that regularly occur between the latitudes of 0° and 45°, which is home to over 75% of the human population [25, 26]. By the second and third days of exposure, we observed increased dam mortality (Supplemental figure 1), 58% of treated dams died during this time frame, and the dams that survived did not have any pups. We terminated the experiment following the institutional animal care recommendation. It was apparent that exposure to 40°C temperatures during the active period for C57Bl6 pregnant mice were lethal to both the dam and the developing fetuses. In all the following experiments, we lowered the exposure temperature to 38°C to prevent maternal and fetal death. 38°C is still considered a heatwave in many latitudes, with many regions experiencing temperatures at 38°C for extended periods of time. In both humans and mice, 38°C induces physiological adaptations to elevated temperatures [27]. Under such experimental model, we found no animals in control and experimental group experienced maternal and fetal mortality.

**Figure 1:**
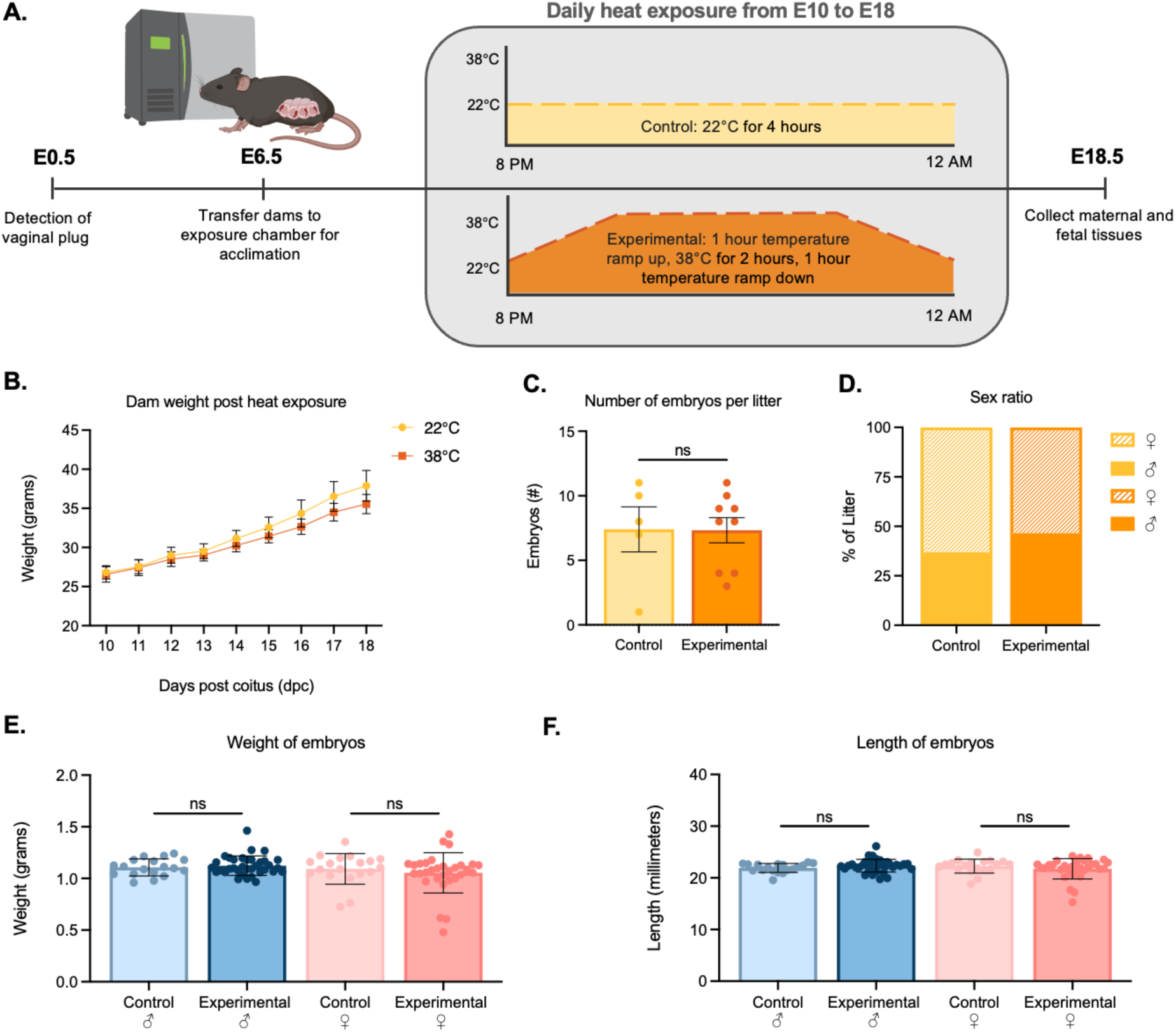
Effects of intermittent gestational heat exposure on maternal weight, litter characteristics, embryo growth, and sex ratio. (A) Schematic of the experimental design. (B) Number of embryos per litter in the control and experimental groups. Each dot represents a single dam. (C) Percentage of male and female embryos within each litter in the control and experimental groups. (D) Maternal body weight over time following heat exposure in the control and experimental groups. (E, F) Embryo weights in the control and experimental groups. Each dot represents a single embryo. For all graphs bars indicate mean ± SEM. Statistical significance was determined using an unpaired t-test. ns, not significant.

### 3.2 Limited impacts of intermittent gestational heat exposure to 38°C on maternal and fetal endpoints

Both control and experimental animals gained similar weight from E10.0-E18.0 (Figure 1B), and had no significant differences in litter size and litter sex ratios (Figure 1C and D). Embryo weights and lengths for both male and female embryos at E18.5 did not significantly differ between the control and experimental groups (Figure 1E). Next, we tested whether stress steroids and sex steroids were altered in response to heat exposure. The levels of corticosterone, aldosterone, testosterone, dihydrotestosterone, estradiol, and progesterone in the sera of experimental animals were not different from those of the control group (Supplemental figure 2). These data indicate that intermittent exposure to 38°C during the second half of gestation does not impact pregnancy outcomes and embryo development.

### 3.3 Placenta morphology and molecular changes are not altered by heat exposure

To determine whether intermittent heat exposure affects placental development, we assessed placental diameter, placental weight, and fetal-to-placental weight ratio at E18.5. In male and female embryos, placental diameter, weight, and fetal-to-placental weight were not significantly different between control and experimental groups (Figure 2A-F). To evaluate whether molecular alterations occurred in the absence of overt structural differences, we performed bulk RNA sequencing on E18.5 placentas. To investigate the determinates of altered male sex differentiation, we focused on male placentas. Principal component analysis (PCA) demonstrated partial separation of samples based on exposure temperature, indicating mild temperature-associated variation in gene expression profiles (Figure 2G). However, differential gene expression analysis (adjusted p < 0.05) identified only five genes between control and experimental placentas, as shown in the volcano plot (Figure 2H). Together, these findings indicate that intermittent gestational exposure to 38°C does not significantly affect gross placental growth parameters in either sex or gene expression profiles in male embryos at E18.5.

**Figure 2:**
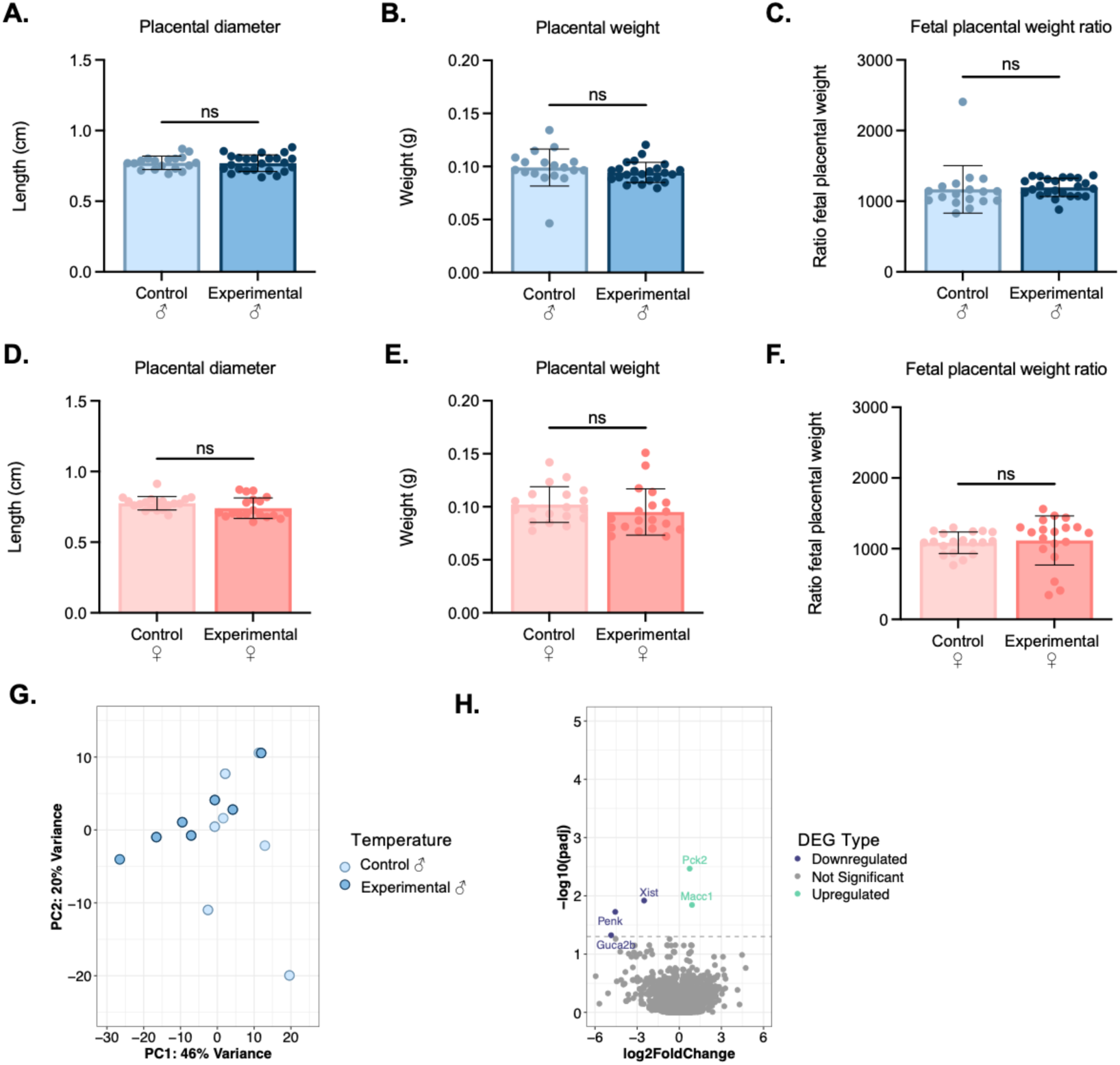
Effects of intermittent gestational heat exposure on placental morphology and transcriptomic profiles in male and female embryos from control and experimental groups. (A–C) Placental diameter (A), placental weight (B), and fetal-to-placental weight ratio (C) in male embryos from the control and experimental groups. Each dot represents a single embryo. (D–F) Placental diameter (D), placental weight (E), and fetal-to-placental weight ratio (F) in female embryos from the control and experimental groups. Each dot represents a single embryo. For all graphs, bars indicate mean ± SEM and statistical significance was determined using an unpaired t-test. ns, not significant. (G) Principal component analysis (PCA) of E18.5 male placental transcriptomes grouped by exposure condition. (H) Volcano plot showing differentially expressed genes (DEGs) between control and experimental male placentas. Differentially expressed genes were defined as adjusted p < 0.05.

### 3.4 Intermittent gestational heat exposure causes mild alterations in differentiation of the external genitalia of the male offspring

Some epidemiological studies have identified negative impacts of heat exposures on external genitalia of male offspring, with boys born in hot climates having higher rates of hypospadias and shorter anogenital distance [10]. Here we only focused on male embryos, as the male genitalia are particularly sensitive to environmental disruptions. To determine if external sex characteristics were altered by intermittent gestational heat exposure, we measured anogenital distance and hypospadias severity of the male embryos at E18.5. We found a significant reduction in anogenital distance in male embryos in the experimental group (Figure 3A-C). After correction for embryo weight and length, anogenital distance remained significantly smaller in the experimental group (Figure 3B and C). In addition to anogenital distance, we quantified the hypospadias severity based on the previously published hypospadias scoring protocol [22]. The control male embryos exhibited a baseline level of mild hypospadias due to individual differences in developmental stage between embryos (Figure 3D). In contrast, hypospadias severity in the experimental group was significantly increased (Figure 3D and E). Four cases of moderate hypospadias that received a score of 2 were found in the experimental, but not the control group.

**Figure 3:**
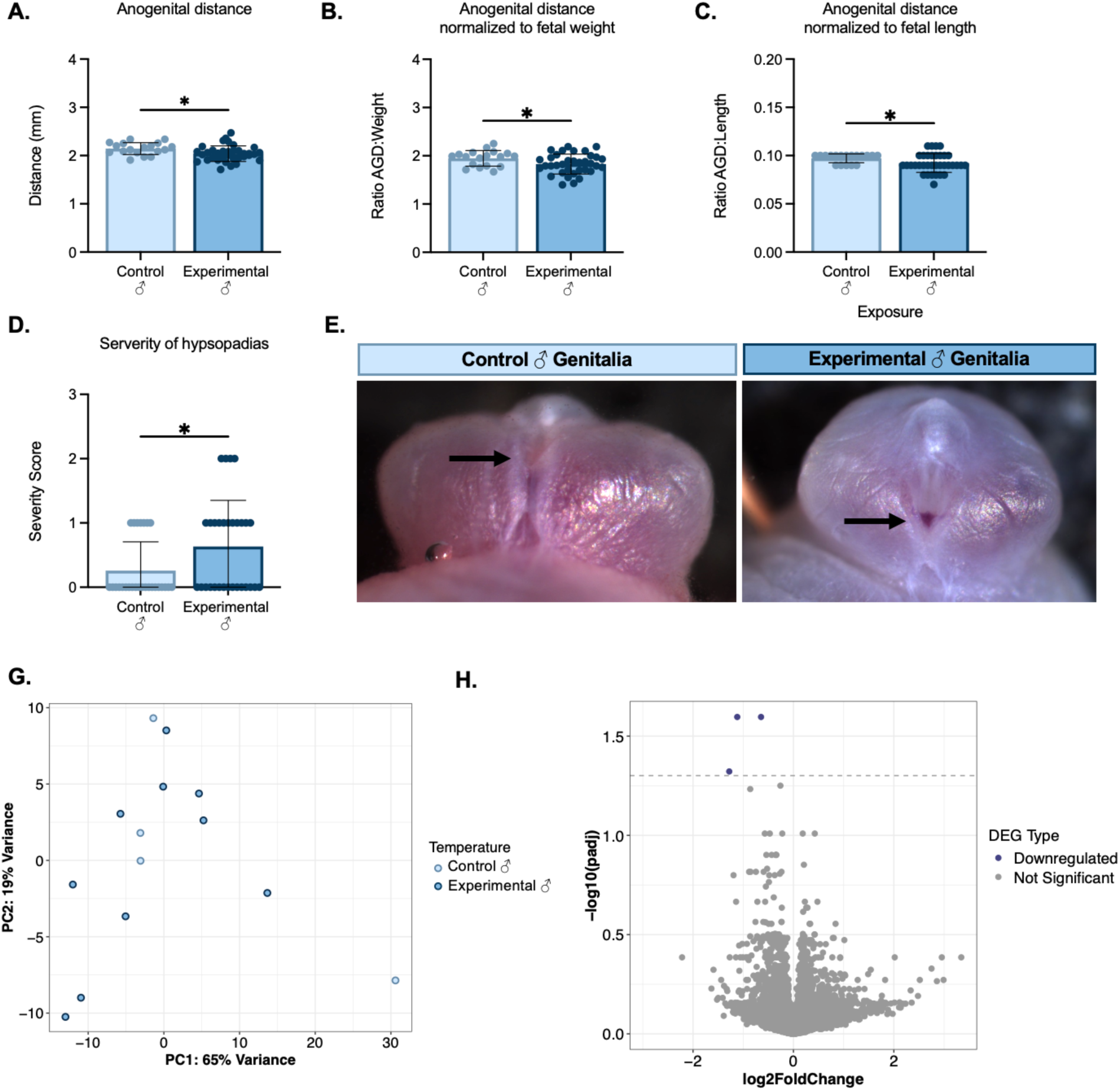
Gestational heat exposure alters male external genital development and gene expression profiles. (A) Anogenital distance in male embryos from the control and experimental groups. Each dot represents a single embryo. (B, C) Anogenital distance normalized to fetal weight (B) or fetal length (C) in male embryos from the control and experimental groups. Each dot represents a single embryo. (D) Objective urethral score evaluation in male embryos from the control and experimental groups. Each dot represents a single embryo. For all graph, bars indicate mean ± SEM and statistical significance was determined using an unpaired Student’s t-test. p < 0.05. (E, F) Representative images of external genitalia from male embryos in the control and experimental groups. (G) Principal component analysis (PCA) of E18.5 male external genitalia transcriptomes grouped by exposure condition. (H) Volcano plot showing differentially expressed genes between control and experimental male external genitalia. Differentially expressed genes were defined as adjusted p < 0.05. (I, J) Gene Ontology biological process enrichment analysis of genes downregulated (I) or upregulated (J) in response to prenatal heat exposure.

To identify the gene expression differences associated with heat induced alterations in penis development, we conducted bulk mRNA sequencing on penises of control and experimental embryos. There was no significant separation in RNA profiles between control and experimental penises based on the PCA analysis axes (Figure 3G). In differential gene expression analysis, 14 genes were significantly upregulated and 29 genes were found to be significantly down regulated in the experimental group (Figure 3H). Gene Ontology (GO) analysis of upregulated genes revealed a significant enrichment of pathways related to angiogenesis, bone remodeling, and transforming growth factor beta stimulus. In contrast, GO analysis of downregulated genes revealed changes in protein catabolic processes, cytoplasmic translation initiation, and a response to Type I interferon. These data show that intermittent gestational heat exposure disrupted external genitalia differentiation in male embryos likely through gene expression changes in developmental and stress response pathways.

### 3.5 Intermittent gestational heat exposure alters the testicular transcriptome at E18.5

Differentiation of male external genitalia requires androgens produced by the fetal testis [11, 12]. To determine whether intermittent gestational heat exposure affects fetal testis development, we performed bulk RNA-sequencing on E18.5 testes from control and experimental embryos. Principal component analysis (PCA) demonstrated separation of samples by exposure temperature, indicating temperature-dependent transcriptional differences (Figure 4A). Differential expression analysis (adjusted p < 0.05) identified 348 significantly altered genes in 38°C exposed testes compared with controls, including 216 upregulated and 132 downregulated genes (Figure 4B). GO analysis of upregulated genes revealed significant enrichment for pathways related to RNA splicing, ribonucleoprotein complex biogenesis, regulation of mRNA metabolic processes, and heterochromatin organization (Figure 4C). Additional enriched terms included regulation of epigenetic gene expression and nuclear RNA splicing, suggesting increased activity in RNA processing and chromatin-associated pathways in response to elevated temperature. As pathways associated with RNA splicing and heterochromatin organization are enriched in developing germ cells, we evaluated germ cell populations. Immunofluorescence staining for Germ cell nuclear antigen (GCNA) showed comparable distribution patterns between control and experimental groups (Figure 4E). Quantification of GCNA-positive cells normalized to testis area revealed no significant difference in germ cell density between groups (unpaired t-test, ns; Figure 4F), consistent with unchanged expression of germ cell genes between groups (Figure 4B).

**Figure 4:**
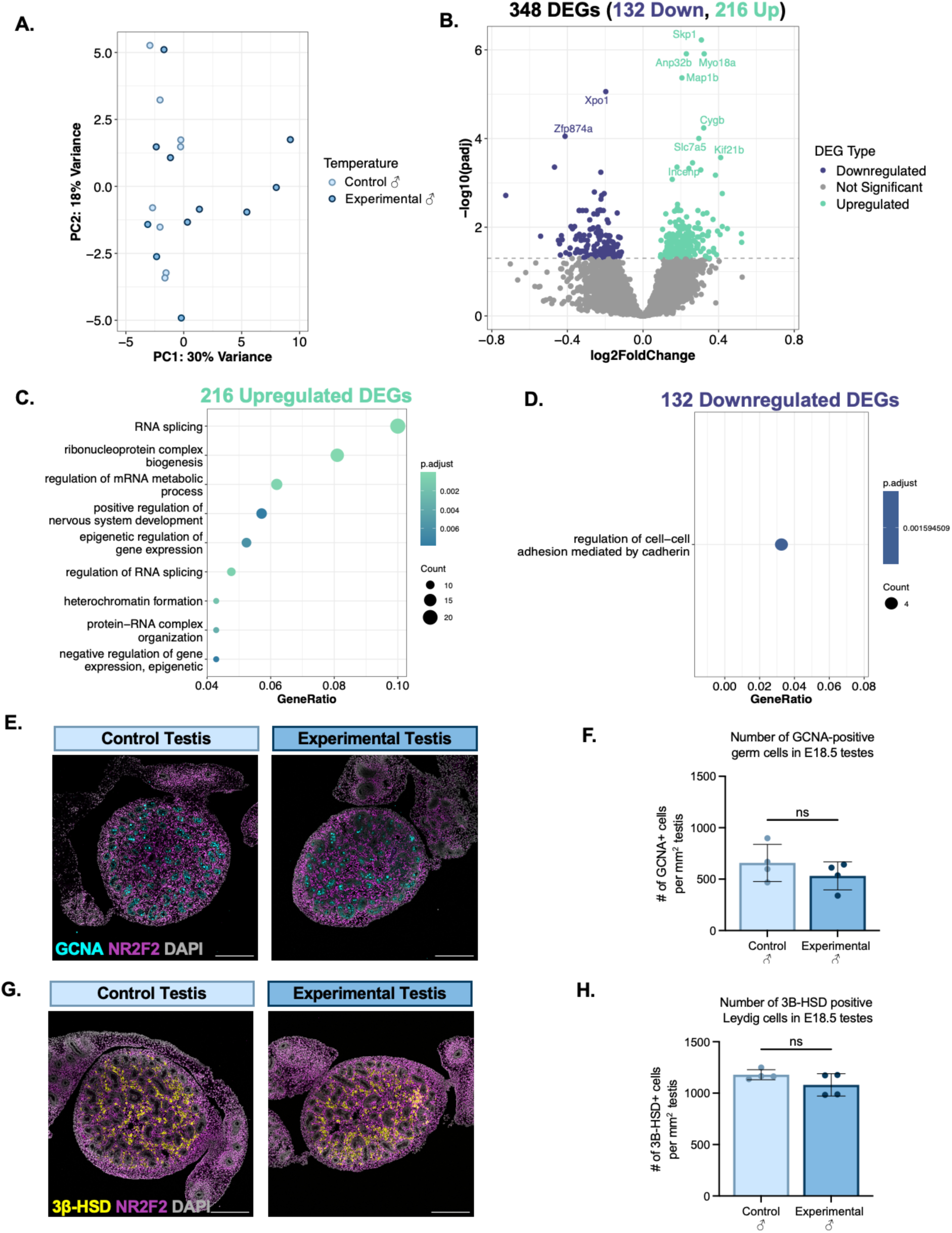
Gestational heat exposure alters gene expression profiles in fetal testes without significantly affecting germ cell or Leydig cell density. (A) Principal component analysis (PCA) of E18.5 testis transcriptomes grouped by exposure condition. (B) Volcano plot showing DEGs between control and experimental testes. DEGs were defined as adjusted p.adj < 0.05. (C, D) Gene Ontology analysis of genes upregulated (C) or downregulated (D) in response to gestational heat exposure. (E) Representative images of E18.5 testes stained for GCNA for germ cells (cyan), NR2F2 for interstitial cells (magenta), and DAPI for nuclear counterstain (gray). Scale bar = 100 µm. (F) Number of GCNA-positive germ cells in E18.5 testes from the control and experimental groups. Each dot represents a single testis section from an individual embryo. (G) Representative images of E18.5 testes stained for 3β-HSD for fetal Leydig cells (yellow), NR2F2 for interstitial cells (magenta), and DAPI for nuclear counterstain (gray). Scale bar = 100 µm. (H) Number of 3β-HSD-positive Leydig cells in E18.5 testes from the control and experimental groups. Each dot represents a single testis section from an individual embryo. For all graphs, bars indicate mean ± SEM. Statistical significance was determined using an unpaired Student’s t-test. ns, not significant.

For the downregulated genes, the GO analysis identified enrichment for regulation of cell–cell adhesion mediated by cadherins (Figure 4D). However, no steroidogenic genes involved in androgen production were altered. This was consistent with the immunostaining of 3β-HSD, a key enzyme in steroidogenesis in fetal Leydig cells. We observed similar localization patterns of 3β-HSD between control and experimental groups (Figure 4G). Quantification of 3β-HSD–positive cells per testis area showed no statistically significant difference between groups (unpaired t-test, ns; Figure 4H).

## 4. Discussion

Pregnant women experience heat stress through a variety of routes, including high fevers, prolonged exposure to elevated ambient heat, and bathing in heated water. In this study, we developed a mouse model of intermittent gestational heat exposure to investigate the effects of elevated environmental temperature on reproductive development of male offspring. Previous studies examining gestational heat stress have primarily exposed animals to heat during their inactive period or throughout the entire day, which may not accurately reflect physiologically relevant environmental heat stress [18, 19, 28, 29]. Our experimental model utilized repeated intermittent heat exposure during the animal’s active period, designed to more closely model cyclical environmental heat stress conditions that may occur during periods of extreme heat. This distinction is important, as the duration, timing, and severity of heat exposure likely influence developmental outcomes and tissue susceptibility.

Our exposure model with daily intermittent 38°C exposure during the second half of gestation did not affect fetal growth or placental development between control and experimental groups, indicating that the observed reproductive phenotypes were not driven by a general growth defect or placental insufficiency. Other heat exposure studies in mice, cow, pig, and sheep have identified significant reductions in placental weight and gene expression, and significant impacts on embryo weight and length [19, 28, 30, 31]. The differences between our study and others are likely attributed to the shorter exposure time in this study. Our study exposed mice for four hours at 38°C, while other studies typically expose to lower temperatures for four hours or longer [18, 19, 28, 29]. Our intermittent exposures may be below the threshold to cause significant differences in placental function and embryo growth. Also, other studies typically put mice directly into a heated cage, as opposed to slowly increasing the temperature with the mouse in the cage. The shock of transferring from room temperature to high ambient heat may induce a stronger stress response in pregnant dams. In human situations, we expect shorter exposures and gradual increases to heat to be more reflective of real scenarios.

We identified abnormalities in male external genital morphology following intermittent gestational heat exposure. To determine whether these phenotypes were associated with altered fetal testis development, we performed transcriptomic analysis of E18.5 testes and identified differential expression of genes involved in RNA splicing, mRNA processing, and chromatin-associated pathways. However, despite transcriptional differences between control and experimental testes, we observed no significant changes in Leydig cell or germ cell abundance, and expression of canonical marker genes for both cell populations remained largely unchanged, suggesting that the external genitalia defects because of heat exposure do not result from defects of androgen production in the fetal testis.

In most adult male mammals, scrotal heat stress causes drastic reductions in testosterone production and overall testis function [32–37]. The reduction in testis-derived androgens then alters androgen signaling in other organs. Typically, the deleterious impacts on the testis are driven by elevated oxidative stress and disruptions in the hypothalamic-pituitary-gonad axis [38–41]. In our data we found no differences in androgen production, Leydig cell number, and steroidogenic gene expression between the control and experimental male embryos. However, we did observe transcriptomic differences in the external genitalia. The experimental group had significant increases in stress related genes, *Psmc4, Psmd1, Ptcd3,* and *Eif31* and decreases in genes associated with differentiation processes, *Sfrp1, Lgr5,* and *Adamts5*. This observation suggests that stress responses in the developing penis could be disruptive to normal penile differentiation and urethra closure. These data together also imply that the impacts of gestational heat exposure on the male reproductive system are androgen independent. Although hypospadias and anogenital distance are thought of as being solely dependent on androgen signaling, <3% human patients with isolated hypospadias have androgen receptor mutations and most patients have normal androgen production [42, 43].

In the testis we did observe extensive gene expression differences related mRNA splicing, mRNA metabolism, and heterochromatin packaging. Some of the differences were attributed to differences in germ cell specific genes, although changes in germ cell numbers were not identified. This demonstrates that the differences in gene expression is not driven by altered germ cell numbers, but by gene expression differences between germ cells. Embryonic germ cells are particularly sensitive to environmental stressors. In several mammalian species, scrotal exposure to heat stress diminishes spermatogenesis capacity, increases sperm abnormalities, and reduces male fertility in adulthood [32, 36]. The limited information in pig offspring suggests that gestational heat stress results in differential genome methylation and reduced sperm production in adulthood [44]. These data further support that gestational heat exposure likely causes epigenetic changes in the testis, particularly the germ cells.

## 5. Conclusion

Our findings demonstrate that intermittent heat exposure during the maternal active period alters reproductive development of male embryos without broadly impairing fetal growth or placental development. Although heat exposure induced abnormalities in external genital morphology and altered fetal testis gene expression, these molecular changes occurred in the absence of major disruptions to androgen-producing Leydig cell or germ cell abundance or expression of canonical cell-type markers. Together, these data suggest that environmentally relevant heat stress may influence male reproductive development through subtle transcriptional and functional mechanisms rather than overt defects in testicular cellular composition. Given the increasing frequency of environmental heat exposure, further studies are needed to determine how gestational heat exposure affects the long-term reproductive health and fertility outcomes.

## Acknowledgements

We thank current and former members of the Yao laboratory for their helpful discussions and technical support. We also acknowledge the NIEHS Comparative Medicine Branch for support with mouse colony maintenance, as well as the Neurobehavioral core facility for technical support and assistance with behavioral studies, and the Epigenomics and DNA Sequencing Core and the Integrative Bioinformatics Support Group for assistance with sequencing and data analysis. This research was supported by the Intramural Research Program (ES102965 to H.H.-C.Y.) of the National Institutes of Health (NIH).

## Author Contributions

K.M.A and C.M.A contributed equally to this work and share first authorship. Conceptualization: C.M.A. and H.H.-C.Y. Methodology: K.M.A, A.K., K.R., B.N., Y.-Y.C., E.O.A, L.A., K.S., J.C., C.G., and H.H.-C.Y. Software: K.M.A, C.M.A, and S.G. Formal analysis: K.M.A, C.M.A, S.G., and H.H.-C.Y. Investigation: K.M.A, A.K., K.R., B.N., Y.- Y.C., E.O.A, L.A., K.S., J.C., C.G., and H.H.-C.Y. Data curation: K.M.A and C.M.A. Writing—original draft: K.M.A and C.M.A. Writing—review and editing: K.M.A, C.M.A, and H.H.-C.Y. Visualization: K.M.A and C.M.A. Supervision: H.H.-C.Y. Project administration: K.R., B.N., and H.H.-C.Y. Funding acquisition: H.H.-C.Y. All authors have read and accepted the data presented in this manuscript.

## Data Availability Statement

The raw data generated and/or analyzed during the current study will be deposited in the Gene Expression Omnibus (GEO) repository and made publicly available upon publication. Custom code used for data processing and analysis will be made available from the corresponding author upon reasonable request.

---

**Supplemental Figure 1:**
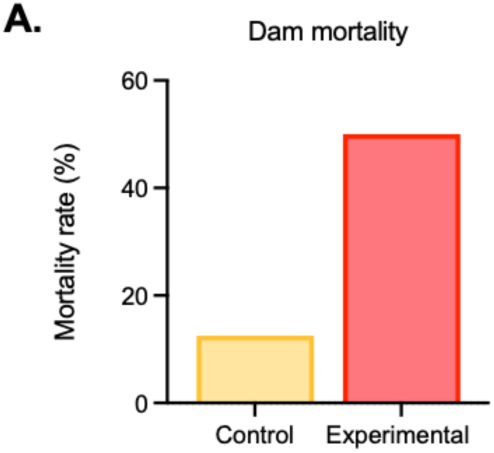
Intermittent gestational heat exposure to 40°C during the second half of pregnancy led to high maternal mortality. (A) Percentage of dams that died during the exposure period.

**Supplemental Figure 2:**
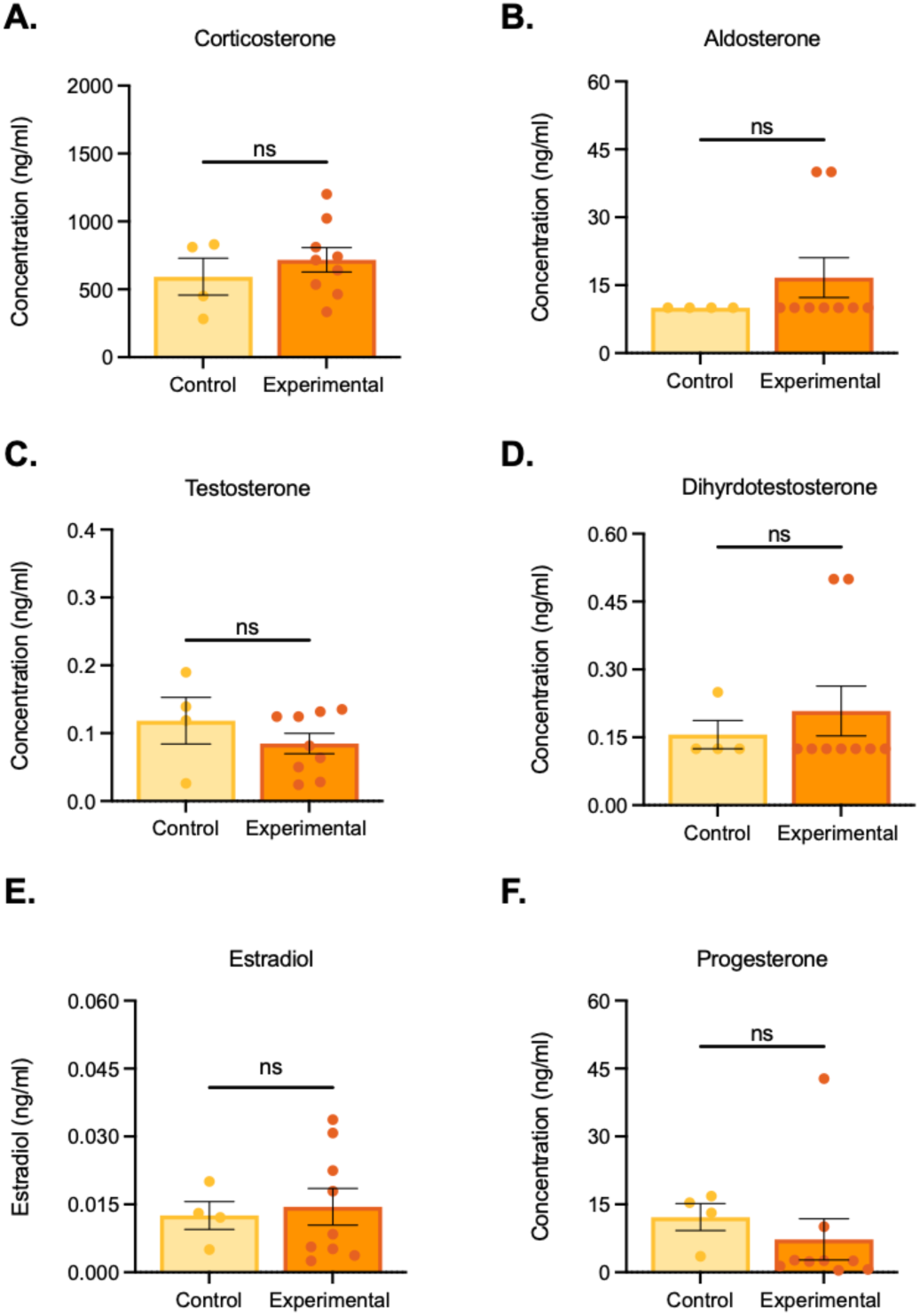
Heat exposure did not alter maternal hormone levels. A-F. Serum levels of corticosterone (A) aldosterone (B) testosterone (C) dihydrotestosterone (D) estradiol (E) or progesterone (F) in dams at E18.5. Each dot represents a single dam. Bar height represents the mean and error bars represent the SEM. Statistical significance was measured using an unpaired t-test. ns, not significant.

